# Quantifying the unobserved protein-coding variants in human populations provides a roadmap for large-scale sequencing projects

**DOI:** 10.1101/030841

**Authors:** James Zou, Gregory Valiant, Paul Valiant, Konrad Karczewski, Siu On Chan, Kaitlin Samocha, Monkol Lek, Exome Aggregation Consortium, Shamil Sunyaev, Mark Daly, Daniel G MacArthur

## Introduction

Recent efforts aggregating the genomes and exomes of tens of thousands of individuals have provided unprecedented insights into the landscape of rare human genetic variation^1,2^ and generated critical resources for clinical and population genetics. The recently announced U.S. Precision Medicine Initiative raises the prospect of growing these databases to encompass hundreds of thousands of human genomes. In the context of these ambitious efforts, it is important to quantify the power of large sequencing projects to discover rare functional genetic variants^3^. In particular, we need to understand, as we sequence ever larger cohorts of individuals, how many new variants we can expect to identify and their expected allele frequencies. Accurate estimates of these quantities will enable better study design and quantitative evaluation of the potential and limitations of these datasets for precision medicine.

## Results

Predicting the number of new variants we expect to identify in larger cohorts requires accurate estimates of allele frequencies of all the genetic variation in the human population, including the rare variants that have not been observed in the current sequencing cohorts^4-6^. The population frequencies of the unobserved rare variants determine the discovery rate of new variants as the cohort sizes increase. We developed a new method, UnseenEst, to estimate the frequency distribution of all variants using the observed site frequency spectrum (SFS) of the current cohort. The method is based on linear program estimators of the SFS^7^, and our mathematical analysis shows that it enables accurate extrapolation of the SFS from current data to cohort sizes more than an order of magnitude larger (Supplementary Information).

Protein-coding variants represent the most readily interpretable and medically relevant slice of human genetic variation, and have been assessed in large sample sizes through the widespread application of exome sequencing approaches^2^. We leveraged data from the Exome Aggregation Consortium (ExAC)^8^ to estimate the discovery rates of different classes of protein coding variants in larger cohorts. We validated UnseenEst by training it on random 10% of the alleles in ExAC and then used the estimated frequency distribution to predict the number of distinct variants that we can identify in the entire ExAC cohort. For every variant type (Supp. Figure 1) and every population (Supp. Figure 2), UnseenEst accurately predicted the number of unique variants that were identified in the entire ExAC cohort as well as the empirical SFS of ExAC (Supp. Table 1).

**Figure 1.**
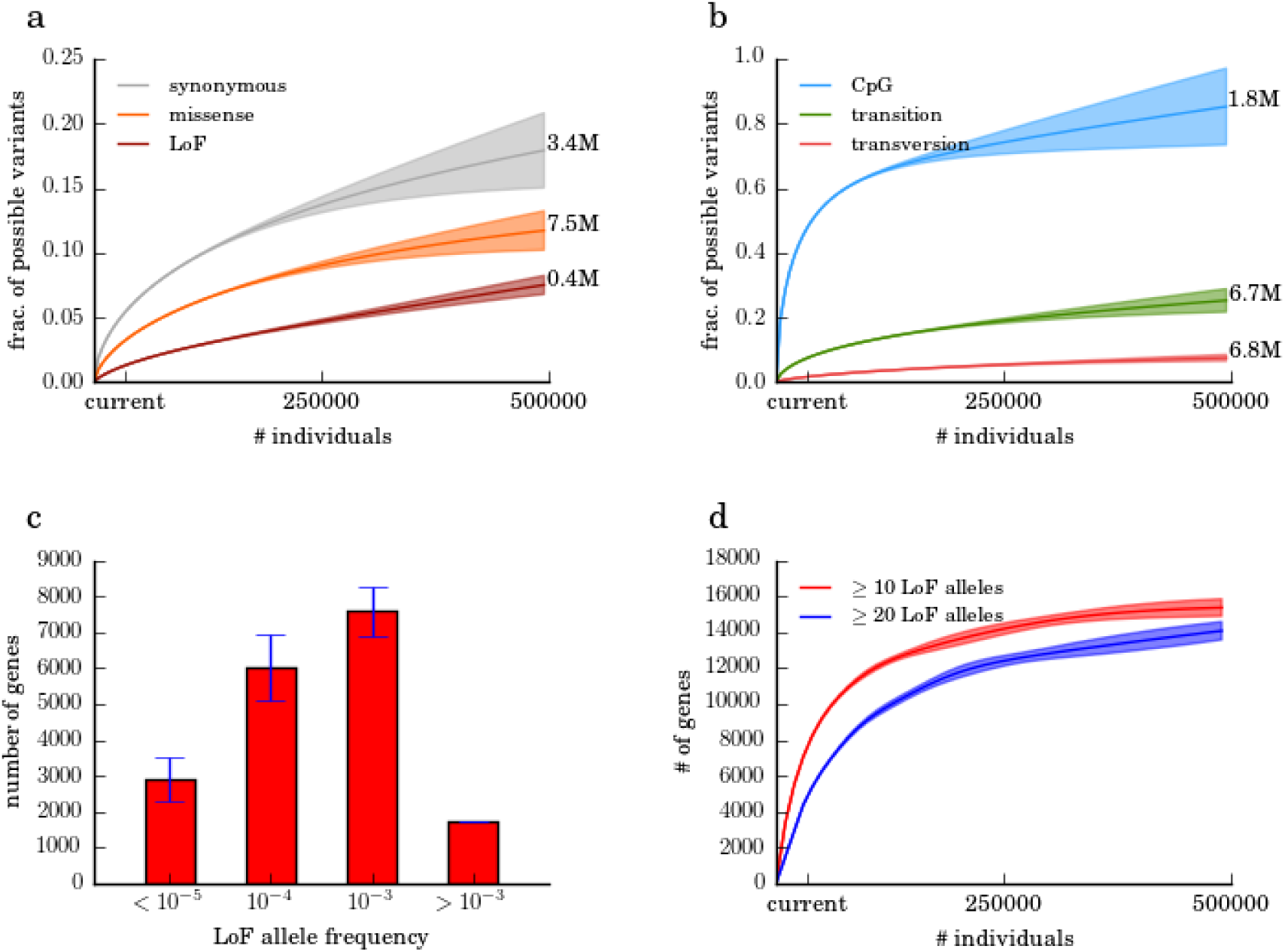
Predictions for the number of unique variants in 500K individuals. We trained UnseenEst on the U.S. Census-matched ExAC cohort (“current”) and predicted the number of unique variants we expect to find in up to 500K individuals. The number of unique variants in the cohort were estimated for synonymous, missense and lose-of-function (LoF) variants in (a), and for CpGs, transitions and transversions in (b). The shaded regions correspond to one standard deviation around the estimates. (c) A gene is classified as LoF on a given allele if that allele contains at least one variant that introduces a stop codon, disrupts a splice donor/receptor site, or disrupts the reading frame. Genes are partitioned into bins based on their LoF allele frequencies: less than 10^−5^, 10^−5^ to 10^−4^, 10^−4^ to 10^−3^, and greater than 10^−3^. The y-axis indicates the number of genes with LoF allele frequency belonging to each bin. Error bars correspond to one standard deviation. (d) Estimated number of genes with at least 10 and 20 LoF alleles.

From the full ExAC dataset, we generated a cohort of 33778 healthy individuals that matched the ancestral population breakdown of the 2010 U.S. Census (Supp. Table 2). We trained UnseenEst on this U.S. Census-matched cohort and predicted the frequency distributions of variants in the entire population (Supp. Figure 3). In particular, we estimated the number of distinct variants we expect to identify in cohorts of up to 500K individuals. These results provide a quantitative framework to evaluate the power and limitations of precision medicine initiatives in discovering rare coding variants.

We categorized the variants by their predicted functional consequence—synonymous, missense, and loss-of-function (LoF), which is defined as point substitutions that introduce stop codons or disrupt splice donor/acceptor sites (Figure 1a). The discovery rate of LoF variants is the lowest, reflecting the fact that LoFs are likely to be deleterious and hence tend to occur comparatively rarely in the healthy population. With 500K individuals, we expect to identify 400K distinct LoF variants or 7.5% of all possible LoF point mutations in the human exome. In the same cohort, we expect to identify 3.4 million synonymous and 7.5 million missense variants, corresponding to 18% and 12% of possible synonymous and missense variants respectively. These estimates indicate that the discovery rates of rare LoF, missense and synonymous variants are far from saturation, even with 500K individuals. We note that slightly higher numbers of distinct synonymous and missense variants (Supp. Figure 4) would be discovered if the 500K individuals were instead sampled from the same ancestral composition as the current ExAC cohort, which contains higher fractions of South and East Asian individuals than the U.S., indicating that the overall discovery rate of rare variants can be boosted by optimizing the population composition of the sequencing cohort.

We additionally classified the variants by their biochemical properties (Figure 1b). With the 34K individuals of the current cohort, we can already identify close to 50% of all possible variants at CpG sites (the most highly mutable substitution class), and the discovery rate for this class of variant quickly saturates as cohorts grow larger. Transversions, in contrast, are discovered much more slowly—attaining 7.6% of all possible transversions with 500K individuals—which is consistent with their much lower mutation rate. We further applied UnseenEst to quantify the number of distinct missense variants we expect to discover in specific gene families of interest, for example genes near GWAS hits and known drug target genes (Supp. Figure 5). Missense mutations in drug target genes are particularly suppressed, suggesting that these genes are more likely to be essential to humans.

LoF variants likely disrupt the normal function of genes and by studying individuals carrying such variants, we can quantify the phenotypic consequence of disrupting particular genes. Therefore, a catalogue of the number of human alleles harboring candidate LoF variants for each gene is an important resource for drug development and disease diagnosis. We applied UnseenEst to estimate the LoF frequency of genes in the U.S. population (Figure 1c, Supp. Figure 6). About 2900 genes have LoF allele frequency lower than 10^−5^, consistent with strong intolerance to inactivation, whereas 1700 genes are expected to harbor LoF variants in at least 0.1% of the population. With 250K individuals, we expect to identify 14K genes that harbor LoFs in at least 10 individuals, substantially expanding the current catalog of 10K such genes in ExAC (Figure 1d, Supp. Figure 7). We estimate that the discovery rate of these genes with multiple LoF occurrences will saturate around 16K, providing an upper bound on the number of genes that can tolerate LoF variants on one allele.

## Discussion

We describe a framework for estimating the power of sequencing cohorts to discover protein-coding variants. We apply it to the largest available collection of sequenced individuals to estimate the discovery power of much larger cohorts such as the ones proposed by the Precision Medicine Initiative. While our predictions here assumed that the samples are representative of the U.S. demography, UnseenEst can be directly applied to estimate the discovery rate of cohorts with different ancestral composition. Our results show that sequencing a cohort of 500K randomly selected U.S. individuals would provide access to over 12% of all possible missense variants and 7.5% of all possible LoF variants, thereby permitting exploration of a substantial fraction of human biological diversity.

---

**Supplementary Figure 1.**
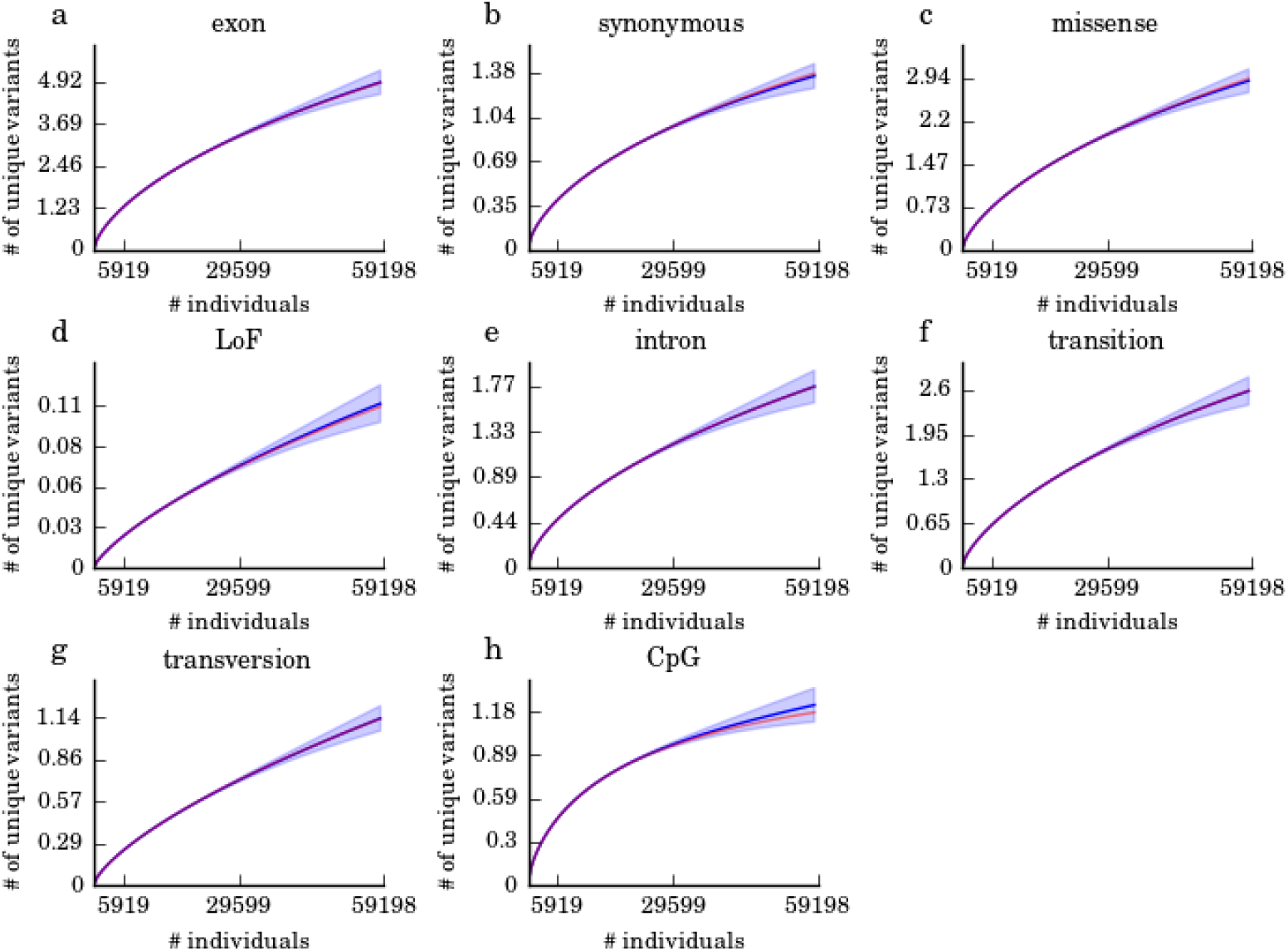
Using 10% of the ExAC alleles to predict the number of unique variants in the entire ExAC cohort. Each panel corresponds to one variant type. For each variant type, we applied UnseenEst on 10% of the ExAC alleles (5919 individuals) to predict the number of unique variants that we would expect to observe in a cohort of size less than or equal to ExAC (59198 individuals). The blue curves are the average predictions over the different 10% sub-samples and the blue shaded regions correspond to one standard deviation from the average. The red curves are the actual number of unique variants observed in ExAC. For all variant types, the predicted number of unique variants is in good agreement with the observed number of unique variants.

**Supplementary Figure 2.**
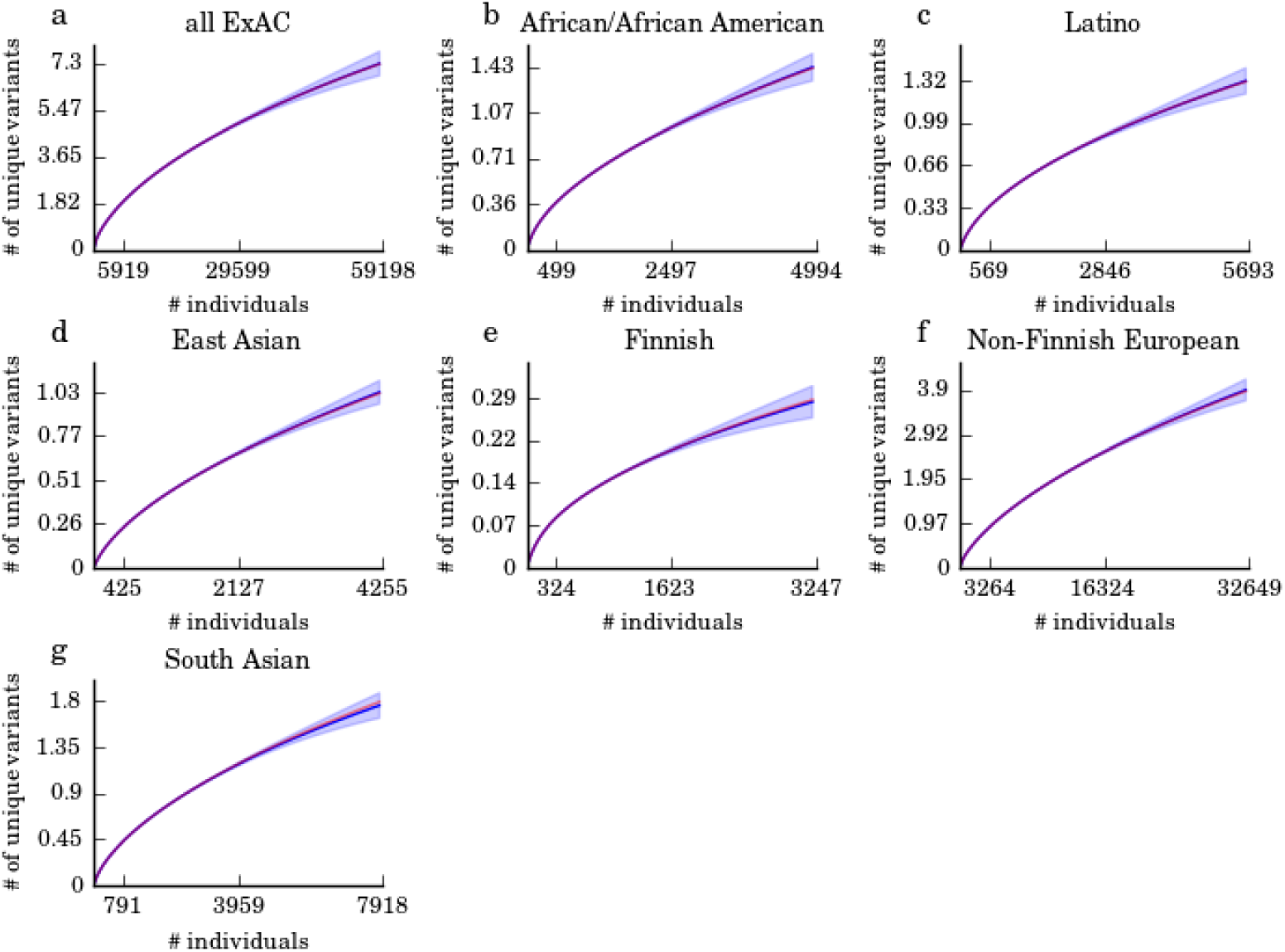
Using 10% of the alleles in each ExAC population to predict the total number of observed variants. For each of the ExAC populations, we trained UnseenEst on random 10% of the alleles and applied it to predict the total number of unique variants in the entire population. The x-axis of each panel indicate the number of individuals of that population; the first mark (e.g. 5919 in (a)) indicate the size of the training set and the last mark (e.g. 59198 in (a)) is the total cohort size of that population in ExAC. The blue curves are the average predictions over the different 10% sub-samples and the blue shaded regions correspond to one standard deviation from the average. The red curves are the actual number of unique variants observed in ExAC. For all variant types, the predicted number of unique variants is in good agreement with the observed number of unique variants.

**Supplementary Figure 3.**
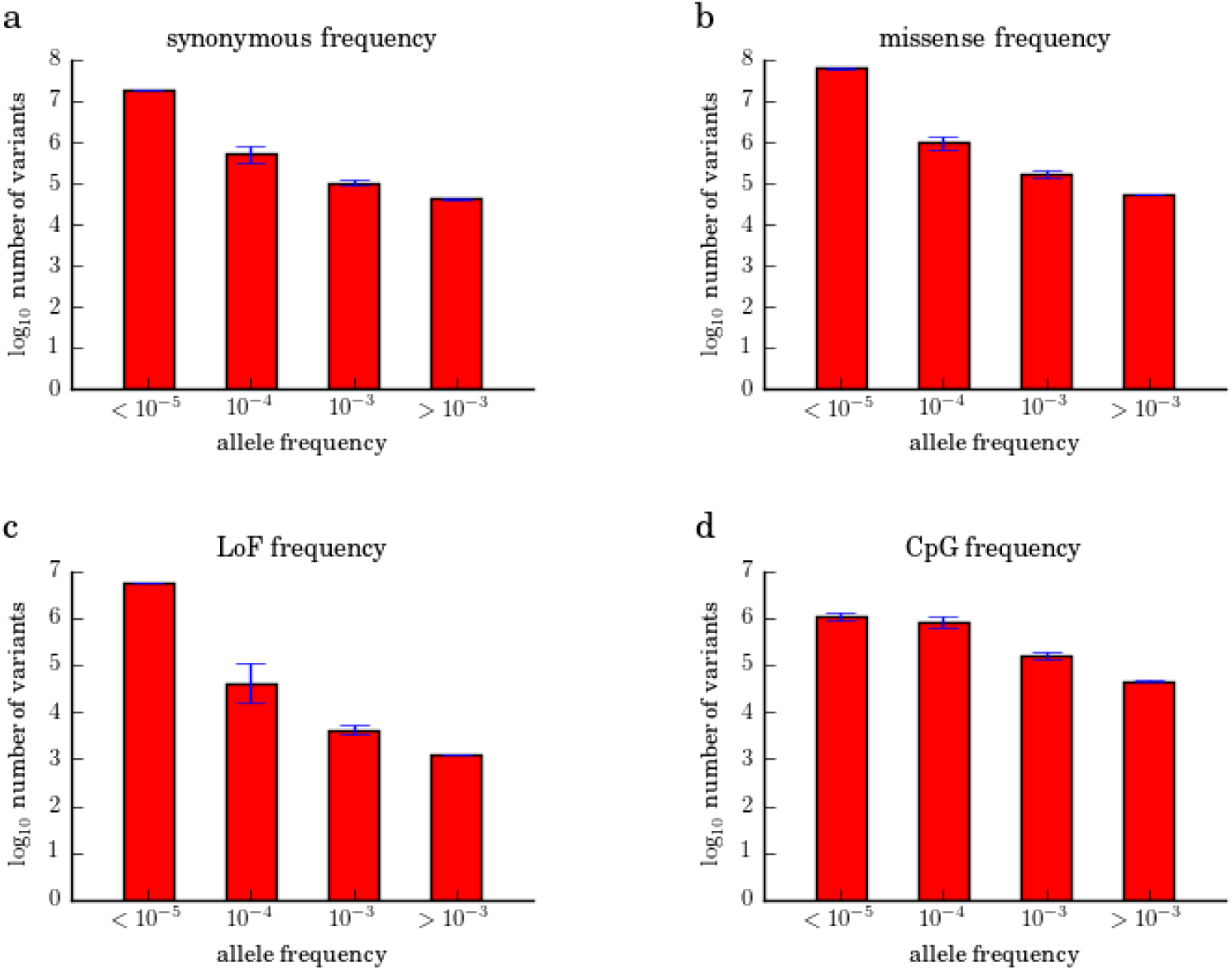
UnseenEst estimated allele frequencies. UnseenEst was trained on the U.S. Census matched ExAC cohort and the synonymous (a), missense (b), LoF (c) and CpG (d) allele frequencies were estimated for the US population. The variants are grouped into bins based on allele frequency: less than 10^−5^, 10^−5^ to 10^−4^, 10^−4^ to 10^−3^, and greater than 10^−3^. The y-axes indicate the log_10_ number of variants in each bin. The error bars correspond to one standard deviation.

**Supplementary Figure 4.**
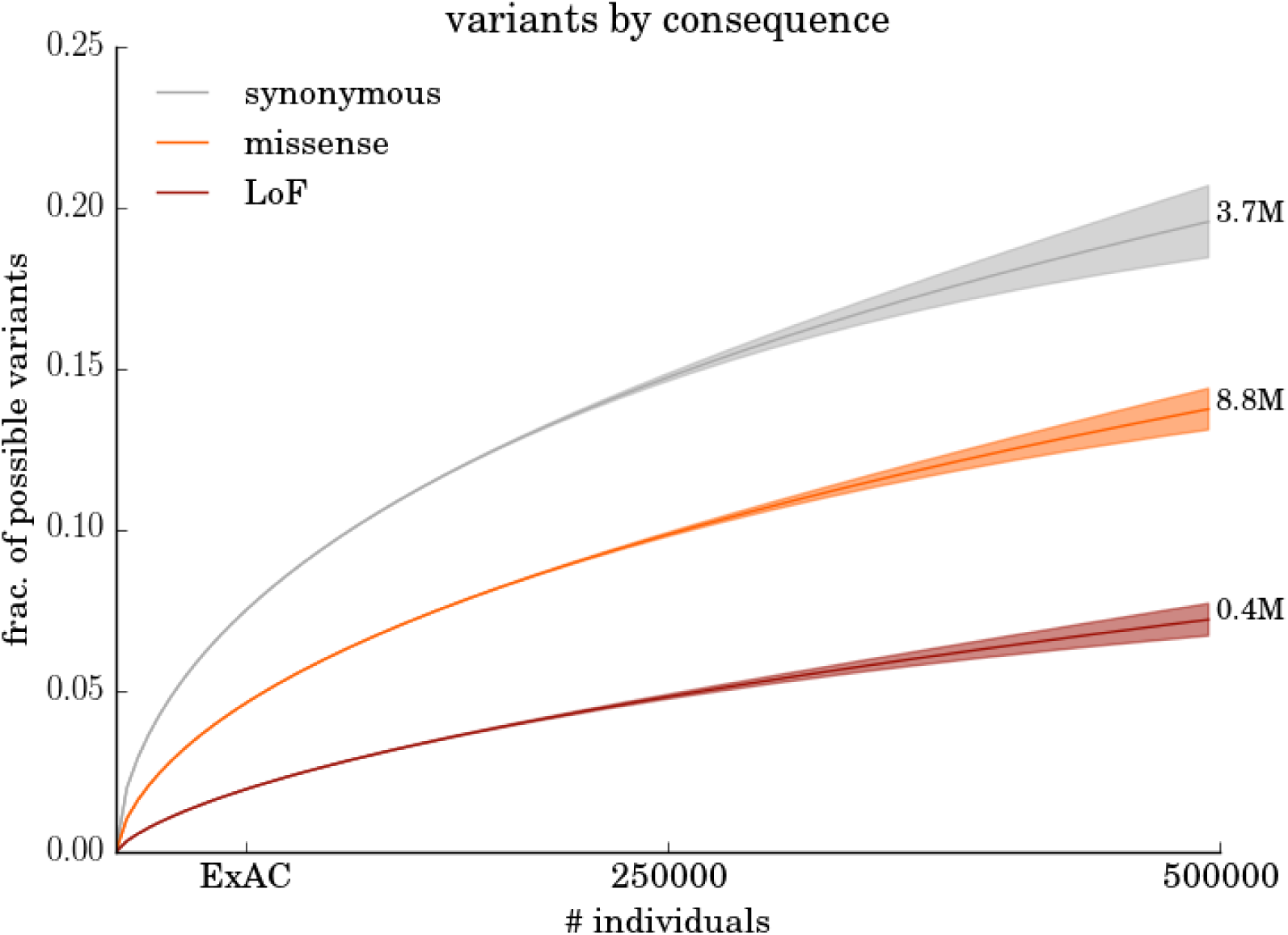
Predicted number of unique variants in cohorts of size up to 500K individuals with the same demographic distribution as the ExAC dataset. The x-axis indicates the number of individuals in the cohort and the y-axis indicates the fraction of possible variants that we expect to observe at in a cohort of that size. We trained UnseenEst on the full ExAC dataset and made the predictions for synonymous (grey), missense (orange) and loss-of-function (brown) variants.

**Supplementary Figure 5.**
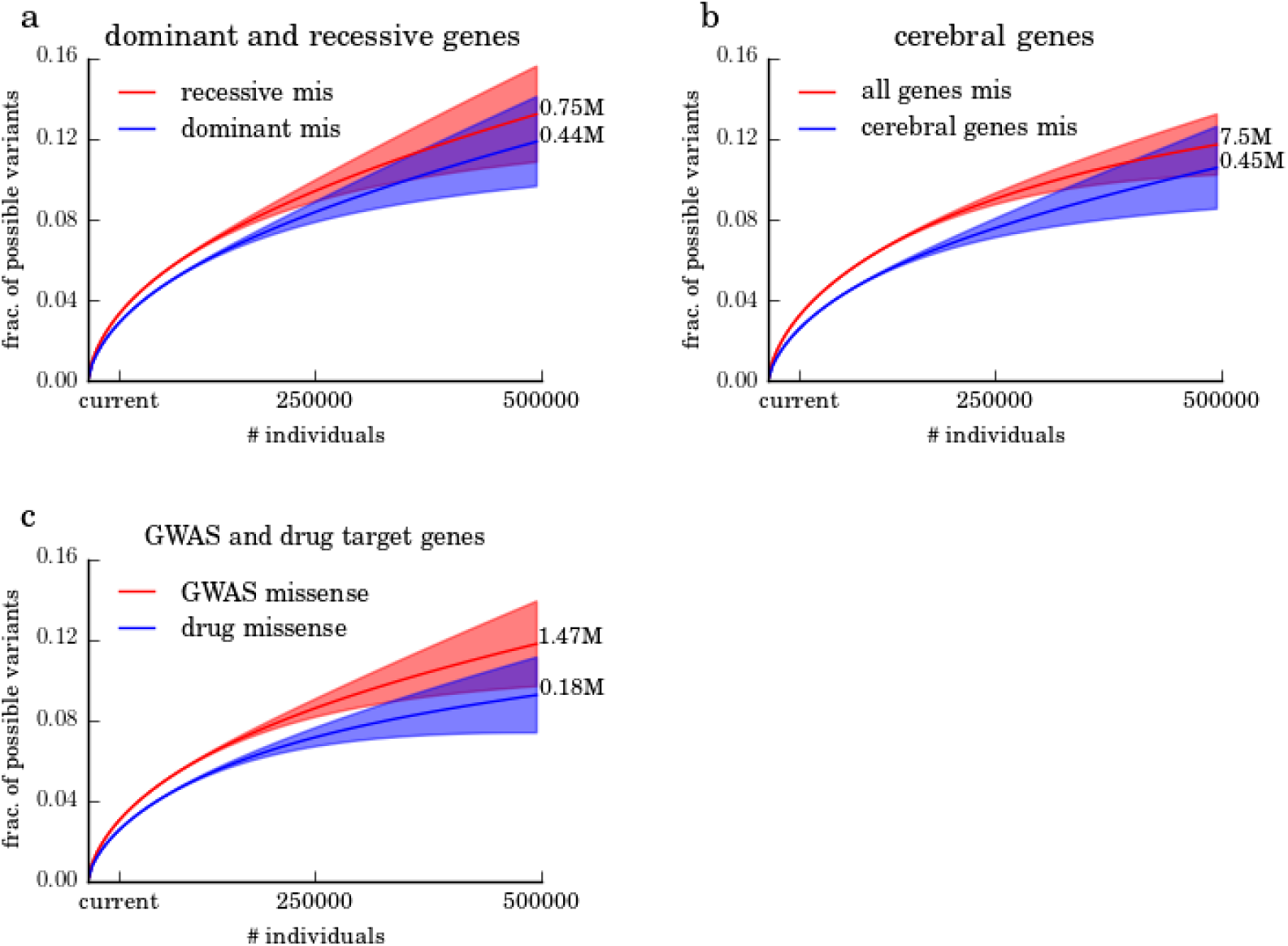
Predicted number of unique missense variants in gene families. We trained the model on the cohort that matches U.S. demographics and predicted the fraction of possible missense variants in each gene family that we can expect to observe in cohorts of size up to 500K individuals. (a) Recessive genes (red) and dominant genes (blue). (b) All genes (red) and genes with cerebral specific expression (blue). (c) Genes associated with GWAS loci (red) and drug target genes (blue).

**Supplementary Figure 6.**
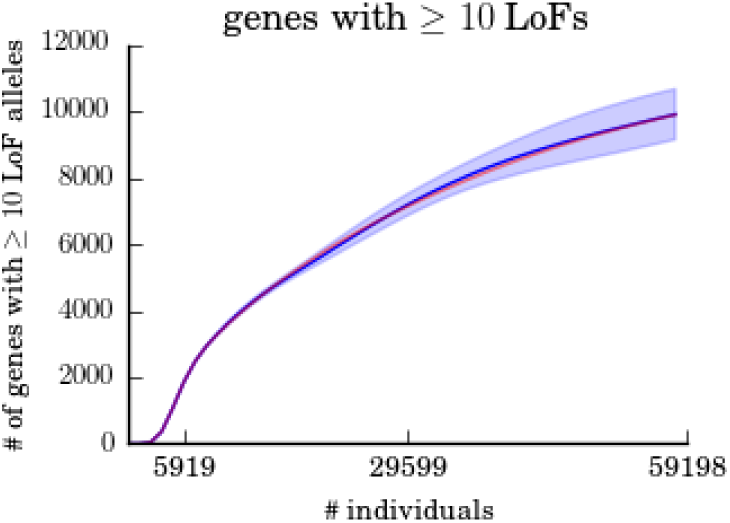
Validation of the estimated number of genes with at least 10 LoF alleles. We trained UnseenEst on random subsamples of 10% of the alleles in the U.S. Census matched cohort and applied it to estimate the number of genes with at least 10 LoF alleles in the entire cohort. The red curve is the actual number of genes with at least 10 LoF alleles and the blue curve is the average predictions over the different subsamples. The shaded blue region corresponds to one standard deviation of the predictions.

**Supplementary Figure 7.**
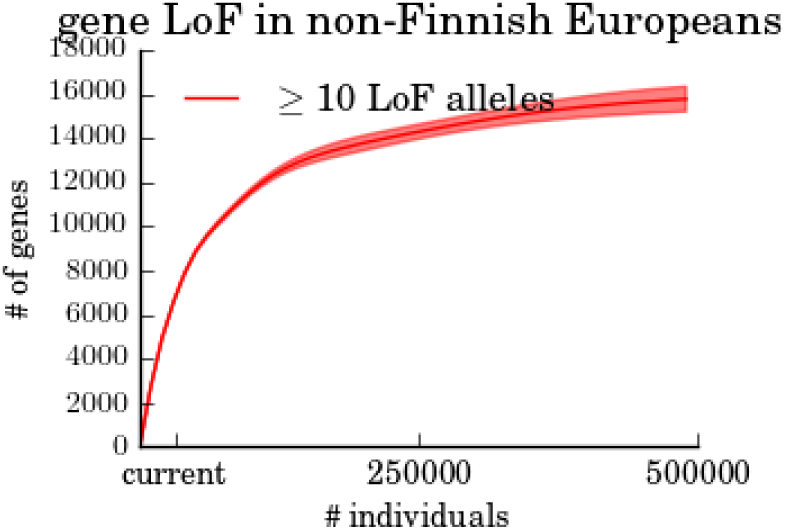
Discovery rate of LoF genes in non-Finnish Europeans. Estimated number of genes with at least 10 LoF alleles in non-Finnish Europeans as a function of the sample size. The number of genes with at least 10 LoF alleles saturates around 16K genes, in agreement with the saturation level of LoF genes in the U.S. Census-matched population (Figure 1d).

**Supplementary Table 1.** Observed and predicted allele counts. Blue rows are the number of ExAC variants with empirical allele counts in bins of 0-10, 10-100, 100-1000, and greater than 1000. Red rows are the predicted allele counts based on UnseenEst trained on 10% of the samples. The standard deviations are shown in the parentheses.

|  | 0-10 | 10-100 | 100-1000 | >1000 |
| --- | --- | --- | --- | --- |
| All variants ExAC | 6.55M | 0.57M | 0.17M | 109422 |
| All variants predicted | 6.59M (0.57M) | 0.55M (0.09M) | 0.18M (0.01M) | 109391 (55) |
| Syn ExAC | 1.22M | 0.13M | 40571 | 27499 |
| Syn predicted | 1.19M (0.12M) | 0.13M (0.02M) | 40658 (2042) | 27446 (187) |
| Mis ExAC | 2.67M | 0.21M | 52794 | 27539 |
| Mis predicted | 2.63M (0.23M) | 0.22M (0.03) | 51957 (2180) | 27352 (178) |
| LoF ExAC | 0.11M | 4300 | 782 | 240 |
| LoF predicted | 0.11M (0.01M) | 4196 (789) | 775 (76) | 225 (12) |

**Supplementary Table 2.** The number of individuals by ancestry. The top row shows the number of individuals of each ancestry in the 2010 U.S. Census. The middle row shows the ancestry composition of the ExAC cohort. The bottom row shows the number of individuals of each ancestry in the ExAC cohort that was adjusted to match the 2010 U.S. Census.

|  | **Non-Hispanic white** | **Latino** | **African-American** | **East Asian** | **South East Asian** | **Total** |
| --- | --- | --- | --- | --- | --- | --- |
| **US Census (2010)** | 196,817,552<br>(65.8%) | 50,477,594<br>(16.9%) | 37,685,848<br>(12.6%) | 10,953,102<br>(3.7%) | 3,374,478<br>(1.1%) | 299,308,574<br>(100%) |
| **ExAC** | 35897<br>(61%) | 5693<br>(9.7%) | 4994<br>(8.5%) | 4255<br>(7.2%) | 7919<br>(13.5%) | 58758<br>(100%) |
| **ExAC census adjusted** | 22212<br>(65.8%) | 5693<br>(16.9%) | 4253<br>(12.6%) | 1249<br>(3.7%) | 371<br>(1.1%) | 33778<br>(100%) |

## Supplementary Note for UnseenEst

### 1 Preliminaries

Given the genetic variation observed in a sample of individuals, what can one infer about all the genetic variation across the entire population? We introduce a robust, general, and theoretically sound algorithm, UnseenEst, for accurately quantifying the distribution of frequencies of all the genetic variation, including the ones that we have not observed in the current samples, based on the sequences from surprisingly small sets of individuals. This estimated distribution of frequencies can then be leveraged to yield accurate estimates of a number of useful properties, including accurate estimates of the number of *new* variants that are likely to be observed in larger cohorts of individuals.

We begin by formalizing the model in which we are working, and describe the sense in which our algorithm recovers the distribution of variant frequencies. The core of our approach is a linear programming (LP) based algorithm, and we discuss the intuition behind this method. We then establish the performance guarantees of our algorithm, proving that, with high probability, it will recover an accurate estimate of the true frequency distribution, and yields accurate predictions for the number of new variants that will be observed in larger samples.

**The model.** Let 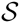 denote a particular variant class of interest. For example, 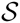 can correspond to all possible missense mutations in a gene family. Each possible variant 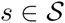 is associated with a probability *p_s_*, which is the probability that an allele contains *s*. We model all the alleles as independent and all variants as independent. Hence the *p_s_*’s are the parameters of independent Bernoulli random variables. When we *sample* an allele, we obtain an independent draw from the Bernoulli at each *s*, 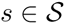. In a sample of *k* alleles, the frequency of observing variant *s* is distributed according to bin(*p_s_*, *k*).

#### Definition 1.1.

*Given* 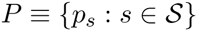*, its histogram is a mapping* 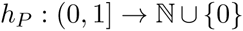, *where*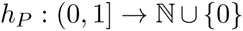 *Informally, h*(*x*) *is the number of variants with probability x. The histogram represents all of the information of P except for the labels of the variants.*

In this work, we are interested in accurately recovering the histogram *h_P_*. For the purpose of estimating any property of the *p_s_*’s that does not depend on the specific labels of the variants themselves, the histogram, *h_P_*, contains all of the useful information. Such properties are referred to as *symmetric* as they are unaffected by ‘renaming’ the variants. The following examples illustrate several interesting symmetric properties:

Examples:
- The total number of variants that occur with probability more than *c* is a symmetric property, and is given by 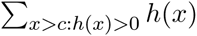.
- The expected number of unique variants that will be observed in a sample of *k* alleles is a symmetric property, and is given by 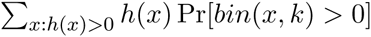.
- The expected number of unique variants that will be observed more than 10 times in a sample of *k* alleles is a symmetric property, and is given by 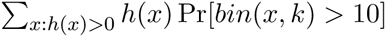.

Because our goal is to recover an accurate approximation of the histogram *h_P_,* it will be useful to define a metric on histograms to provide a concrete notion of what it means for two histograms to be “similar”.

#### Definition 1.2.

*Given two histograms, g and h, assume without loss of generality that* 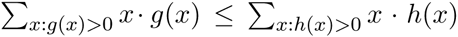. *The* generalized relative earthmover distance *between them, denoted R*(*g, h*), *is defined to be* 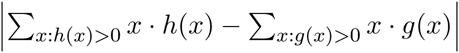 plus the minimum over all schemes *of moving the mass of histogram g to yield h*′*, where*

- *h′ is any histogram such that* 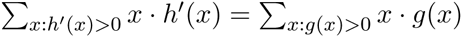 *and h′*(*x*) ≤ *h*(*x*) ∀*x;*
- *the cost, per unit mass, of moving from probability value x to probability y is* 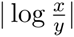.

*Note that the amount of mass in histogram g at probability value x is given by x · g*(*x*).

The following example illustrates this definition.

#### Example 1.3.

*Let h denote the histogram representing* 200 *variants that each occur with probability* 1/100. *Hence h*(1/100) = 200*, and for all x ≠* 1/100, *h*(*x*) = 0*. Let g denote the histogram consisting of* 50 *variants with probability* 1/100, *and* 300 *variants that occur with probability* 1/200, *hence g*(1/100) = 50, *and g*(1/200) = 300. *Note that both histograms have the same total mass, since* 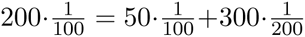*. The relative earthmover distance satisfies* 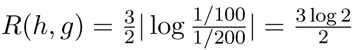*, since g can be obtained from h by moving* 3/2 *mass from probability* 1/100 *to probability* 1/200 *to yield histogram g.*

The generalized relative earthmover distance allows for comparisons of histograms with different total masses, which is necessary since the inferred histogram from data will typically have a slightly different mass from the true distribution. Intuitively, relative earthmover also highlights the importance of estimating the rare variants well: mistaking variants with frequency 10^−5^ for frequency 10^−6^ suffers substantial distance cost. The other main reason for using the relative earthmover distance is that many properties of interest are Lipschitz continuous with respect to this distance: if two histograms are close in relative earthmover distance, then they have similar property values. In particular, if we guarantee that, with high probability, our algorithm recovers an estimate of the underlying histogram that is accurate in relative earthmover distance, then estimates of properties that we obtain from the recovered histogram will be accurate.

The following proposition, whose proof is given in Section 6.3 illustrates this point, and shows that if two histograms are close in relative earthmover distance, then the expected number of variants that will be observed in any given sized sample will be correspondingly similar.

#### Proposition 1.4.

*Given two lists of probabilities* 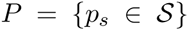 *and* 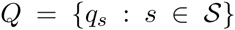, *let* 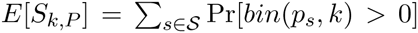 *denote the expected number of variants observed in a sample of k alleles with the distribution of frequencies given by P, and let E*[*S_k,Q_*] *denote the analogous quantity corresponding to frequencies Q. Then, for any k* > 3,

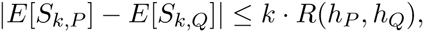

*where R*(*h_P_, h_Q_*) *is the generalized relative earthmover distance between the histograms corresponding to P and Q.*

In analogy to the histogram *h_P_* giving us a label-less representation of the true underlying *P* = {*p_s_*}, it is convenient to have a label-less representation of the observed variant counts from a sample of alleles. To this end, we define the *fingerprint* of the observed variants, which is also known as the site frequency spectrum (SFS) in genetics, or the “pattern” of the sample in some statistics contexts.

Definition 1.5.

*Given sample X of k alleles, the associated **fingerprint,*** 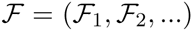 *is the “histogram of the histogram” of X. Formally,* 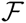 *is the vector whose ith component,* 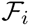*, is the number of variants in* 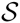 *that occur exactly i times in sample X.*

**Remarks on the model.** Our model assumes that all the variants are independent random variables. Population demography and linkage disequilibrium introduce correlations especially between the common genetic variants. For the common variants, UnseenEst uses the empirical frequency to accurately estimate the true population frequency. For the very rare variants, which UnseenEst tries to estimate while using the independence assumption, this assumption is also a better approximation of the real data.

While the discussion here focuses on estimating the histogram of genetic variation, UnseenEst is an general approach to estimate the histogram and statistical properties of any finite lists of probabilities {*p*_1_,…,*p_n_*} from independent Bernoulli samples, and can have broad applications beyond genetics. Note that 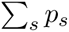 can be significantly smaller or larger than 1.

### 2 The UnseenEst Algorithm

We partion the variants into two classes: *common* variants and *rare* variants. In our applications, variants with empirical allele frequency above 1% are defined to be common. With current cohorts of 10s of thousands of alleles, we are likely to have observed all the common variants and the empirical allele frequencies of the common variants should be very close to the true population frequencies. Therefore we focus the efforts of the algorithm on estimating the frequencies of rare variants.

Given a sample of *k* alleles and the associated fingerprint 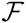, we truncate the fingerprint to only the rare variants with frequency less than 1%, i.e. we consider 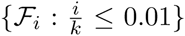. For the common variants with frequency above 1%, we simply use their empirical frequency as an estimator of the true frequency. On the truncated fingerprint, we solve the following linear program for variables corresponding to *h*(*x*), *x* ∈ *X* for a finite mesh of probabilities *X*.

#### Algorithm UnseenEst.

**Input:**

- Fingerprint 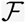 from k alleles.
- A set of probability values 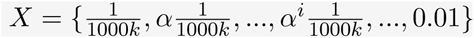. We use *α* = 1.05.
- *n* = upper bound on the number of possible variants.

**Output:** histogram 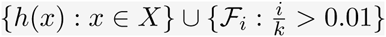.

Solve for *h*(*x*), *x* ∈ *X*, to minimize the objective function

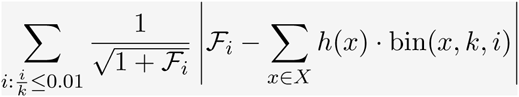

subject to the constraints

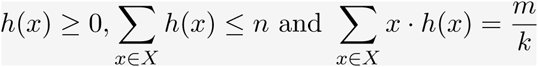

where 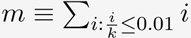 is the total number of observed variants with empirical frequency less than or equal to 1%.

For a given histogram *h*, 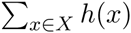 is the expected number of variants observed *i* times in *k* alleles. The objective function of the LP captures how much this expected number of variants deviates from the empirical number of variants observed *i* times (represented by the entries of the fingerprint 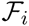). The term 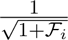 normalizes the deviation by the standard deviation. The constraints enforce that *h*(*x*) ≥ 0, namely that there can not be a negative number of variants that arise with a given probability, and that the total sum of the probabilities matches the empirical estimate of the sum of the probabilities. Note that 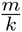 is a very accurate estimator of ∑ *p_s_* for large *k*. In practice, we found it sufficient to use the probability mesh *X* with geometrically increasing probabilities with rate *α* = 1.05. Using a smaller *α* can marginally improve accuracy at the cost of run-time. UnseenEst is very efficient; on the ExAC dataset (described below) the computation took less than 10s on a standard laptop.

**Estimating the number of unique variants.** Given the estimated histogram *h* produced by UnseenEst, the expected number of unique variants in a sample of *k* alleles is

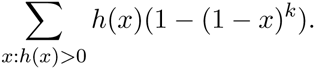

#### 2.1 Performance Guarantees

For the performance guarantees, we analyze the slightly modified linear program below. To simplify the notations, we set the constants *B, C, D* such that

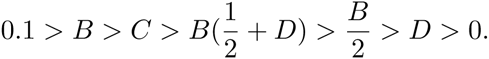

Given as input an untruncated fingerprint 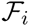 of *m* total variants generated from *k* alleles, the linear program algorithm is

Algorithm UnseenEst2.

**Input:** fingerprint 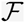 from *k* alleles, *n* = upper bound on the number of possible variants, and 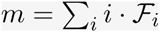 is the total number of variants observed in *k* alleles.

- Define the set 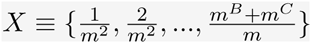.
- For each *x* ∈ *X*, define the associated LP variable *h*(*x*).

**Output:** histogram *h* with support on *X*.

Solve for *h*(*x*), *x* ∈ *X*, to minimize the objective function

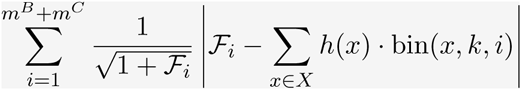

subject to the constraints

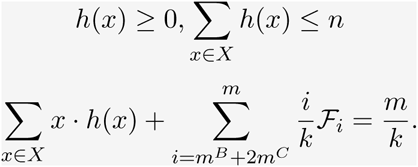

For each integer *j* ≥ *m^B^* + 2*m^C^*, set 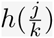 to 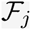.

The UnseenEst2 algorithm satisfies the following guarantee.

##### Theorem 2.1.

*Let n be the support size (the number of possible variants), k be the number of alleles sequenced, and P* = {*p_s_*} *denote the true distribution of the variant frequencies with* ∑*p_s_ the expected number of variants per allele. For sufficiently large n, with probability at least* 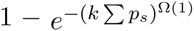, *the algorithm will return a histogram g satisfying:*

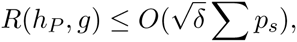

*where* 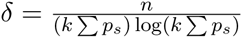 *and the ‘O’ notation hides an absolute constant.*

The above theorem, together with Proposition 1.4 implies the following corollary:

##### Corollary 2.2.

*Let n be the support size (the number of possible variants), k be the number of alleles sequenced and* ∑ *p_s_ is the expected number of variants per allele. Given a sample of k alleles, with probability at least* 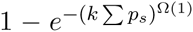*, the algorithm estimates the expected number of unique variants that will be observed in a sample of k′ alleles to within additive error*

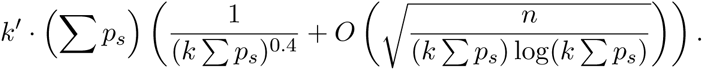

One interpretation of the above corollary is that the estimate of the expected number of unique variants will be accurate, relative to the total expected number of observed variants, *k*′ ∑ *p_s_*, provided *n* < (*k* ∑*p_s_*) log(*k* ∑*p_s_*). For comparison, the naive algorithm that attempts to learn the distribution P will only be accurate in the regime where *n* < *k* ∑*p_s_*.

### 3 Datasets

We used the exome sequencing data from the Exome Aggregation Consortium (ExAC) [1]. This dataset consists of high-quality sequencing of the protein-coding regions in the genome (exomes) from 60706 healthy individuals. Consistent with the ExAC analysis, we considered only regions of the exome with sufficient sequencing depth: each nucleotide must be covered by at least 10 reads in at least 80% of all ExAC individuals.

**Loss-of-function (LoF) variants.** We define LoF variants to be single-nucleotide substitutions that introduce a stop codon in the reading frame or disrupts a splice donor or receptor site. We do not include insertion/deletions (indels) in the class of LoF variants. Variant annotation was performed using the Variant Effect Predictor (VEP) v81 on Gencode v19 and genome build GRch37. LoF annotation was performed using LOFTEE (version 0.2; available at available at https://github.com/konradjk/loftee) plugin to VEP. While early stop codon and splice donor/receptor disruptions often lead to truncated proteins, this does not imply that the protein has lost all of its function. Our annotation of LoF variants does not explicitly assess protein function and hence serves only as a proxy for the true deleteriousness of the variant.

**Upper bound on the number of possible variants.** A natural way to interpret the discovery rate of a given variant class is to calculate, among all possible variants in this class, what fraction of them do we expect to observe at a given sample size. To estimate an upper bound for the total number of possible variants in each class, we first identified all the nucleotides for which we have sufficient read coverage (at least 10 reads in at least 80% of all ExAC individuals). Then at each well-covered nucleotide we identified the number of possible variants that belongs to a given class. For example, if the reference genome at a particular nucleotide is *A*, then there are two possible transversions (*A* → *C* and *A* → *T*) and one possible transition (*A* → *G*). The upper bound for the number of possible transversions is then calculated as the sum of the possible transversions across all well-covered nucleotides (which is just 2 times the number of well-covered nucleotides), and similarly for other variant classes. The upper bound on the number of possible variants in each variant class in the ExAC data is given below.

| Variant class | Upper bound on the # of possible variants |
| --- | --- |
| LoF | 5,720,461 |
| CpG | 2,086,001 |
| synonymous | 18,762,312 |
| missense | 63,986,829 |
| missense in cerebral genes | 4,204,277 |
| missense in dominant genes | 3,678,497 |
| missense in drug target genes | 1,908,788 |
| missense in GWAS genes | 12,517,761 |
| missense in recessive genes | 5,628,661 |
| transitions | 45,824,366 |
| transversion | 91,648,732 |

For each variant class, we divided the number of unique variants we expect to identify by this upper bound to obtained the *fraction of possible variants* observed at a given cohort size. As technology improves in future sequencing projects, we expect the well-covered regions of the exome to increase and hence the number of identified variants to also increase.

**LoF genes.**

We used the same set of 18225 genes as in the ExAC analysis [1]. Briefly, we summed all exon level variant counts across Gencode v.19 canonical transcripts. If an exon had a median depth < 1, the variant counts for that exon were not included in the total for the transcript. We then removed all transcripts where no variants were observed. We also removed the outliers whose observed synonymous and missense counts deviated significantly from the expected. This left 18225 for which ExAC had high-quality data.

We associated with each gene, *g*, a Bernoulli random variable with probability *p_g_*, which corresponds to the probability that an allele of the gene contains at least one LoF variant as defined above or at least one insertion-deletion (indel) that disrupts the reading frame. The presence of such a LoF variant or indel is a proxy for true loss-of-function and does not necessarily mean that the gene is entirely non-functional on that allele. For example, if the LoF variant introduces a stop codon near the 3’ end of the gene, then the corresponding truncated protein may still retain some functions.

UnseenEst can be applied to estimate the histogram of any set of probabilities {*p_g_*}, and hence it directly applies in this setting. On the U.S. Census matched cohort, we assign a gene 1 on an allele if it has at least one LoF variant or frame-shift indel. Otherwise it is assigned a 0. The fingerprint 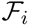 here corresponds to the number of genes that are LoF in exactly *i* alleles. We trained UnseenEst on this gene-level fingerprint.

**Gene lists.** We describe the curation of the various gene lists below.

- **Dominant genes:** 691 OMIM disease genes deemed to follow autosomal dominant inheritance according to [2][3].
- **Recessive genes:** 1163 OMIM disease genes deemed to follow autosomal recessive inheritance according to [2][3].
- **GWAS genes:** 2801 genes that are the closest 3’ and 5’ genes to GWAS hits in the NHGRI GWAS catalog as of February 9, 2015.
- **Drug target genes:** 460 genes whose protein products are known to be the mechanistic targets of drugs; curated from [4][5].
- **Genes with cerebral specific expression:** 979 genes with cerebral specific expression downloaded from [6].

### 4 Validation experiments

We performed multiple experiments to validate the prediction accuracy of UnseenEst.

**Accuracy of allele frequency estimation.** For each class of variant (synonymous, missense, LoF, CpG) we randomly partitioned all the ExAC alleles into ten groups. We trained UnseenEst on the site frequency spectrum of one partition (i.e. 10% of the alleles) and used the model to predict the allele frequency distribution of the entire ExAC cohort. We grouped variants into 4 frequency bins: 1) variants that occur in 0–10 alleles; 2) variants that occur in 11–100 alleles; 3) variants that occur in 101–1000 alleles; and 4) variants that occur in more than 1000 alleles. We repeated this procedure for each of the ten random partitions and computed the average and the standard deviation for the number of variants predicted to belong to each bin. These estimates are compared with the observed number of variants in each bin in ExAC.

**Accuracy of the estimated number of unique variants.** For each variant class, we randomly sampled 10% of the alleles and applied UnseenEst on the SFS of this subsample to estimate the histogram 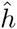 of the variant frequencies. For any positive integer *k*, the number fo unique variants we expect to see in *k* alleles is ∑*_p_ h*(*p*)(1 − (1 − *p*)*^k^*). As before, we compute the average and the standard deviation of the estimates across the different 10% subsamples. To produce the ‘true’ discovery rate, we create a random ordering of all the ExAC alleles. Then for each *k* less than the ExAC cohort size, we count the number of unique variants observed in the first *k* alleles.

**Accuracy of gene LoF frequency.** We randomly partitioned the alleles into ten subsets. For each subset with 10% of the alleles, we generated the gene-level LoF fingerprint from this subsample. We trained UnseenEst on this subsampled fingerprint and compared the predicted number of genes with at least 10 LoF alleles with that of the observed in the entire ExAC data. The mean and standard deviations of the predictions were computed from the 10 different partitions.

### 5 Related works

While our algorithm and analysis are closely related to the approach in [7][8], there are important differences in the model. In [7], we have an unknown discrete distribution *P* on *n* elements and we have *k* independent samples from *P*. This model was motivated by the classic problem of estimating the vocabulary size of Shakespeare from a sample of his works [9]. The discrete distribution setting can be reformulated by associating with each element *s* an independent Poisson random variable poi(*p_s_*), where *p_s_* is the weight of *P* for *s*. Here, unlike in our model, ∑*_s_p_s_* = 1. The number of times that *s* appears in *k* samples is distributed according to poi(*k* · *p_s_*). In our genetics model, the number of times that a variant *s* appears in *k* alleles is distributed according to bin(*p_s_*, *k*). While poi(*k* · *p_s_*) and bin(*p_s_*, *k*) both have expectation *k* · *p_s_*, the Poisson has a slightly larger variance. Because the number of elements *n* is potentially very large, this difference between Poisson and Binomial aggregates over all the elements and can give rise to substantial differences in the expected fingerprints between the two models.

Recently, [10] also proposed using linear program to estimate the discovery rate of new variants. They solve two linear programs with hypergeometric coefficients, to estimate the upper and lower bounds on the number of unique variants at a given sample size that are consistent with the observed site frequency spectrum. Under the infinite genome assumption (i.e. there are infinitely many possible variants), [10] showed that there exist solutions to these two linear programs. The approach of [10] tries to identify the range of the number of unique variants that is consistent with the observed data, though it does not guarantee how wide this interval is and whether it concentrates around the true value in general. Our linear program is guaranteed to produce a histogram that is close to the true SFS. Moreover our analysis makes explicit the dependence on the sample size *k* and the frequency distribution *p_s_* which was not present in [10].

Bayesian approaches have also been applied to estimate the number of unseen variants [11] [12]. Other approaches based on the jackknife estimator have also been applied to similar settings [13][4]. In [11], the mutation probabilities of the variants are assumed to be i.i.d. samples from a Beta(*a, b*) prior, where the hyperparameters *a, b* are fitted from data. A limitation of this approach is that it requires parametric forms for the distribution of variant frequencies, which requires some model of demography and selection. For example, the Beta prior used in [11] is a reasonable assumption for neutrally evolving variants but may not be appropriate for deleterious mutations. The advantage of UnseenEst is that it does not require any modeling assumption about selective pressure and demographic history, i.e. it is non-parametric. Theorem 2.1 applies in all settings where the independence assumption is a reasonable approximation.

### 6 Proofs of the Guarantees

The proof of Theorem 2.1 for UnseenEst2 has three main components. First we show that given a sample of *k* alleles from the above model, with high probability the empirical fingerprint 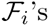 are close to their expected values 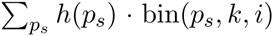. This sample of *k* alleles is what we call a *faithful* sample. Next we show that given a faithful sample, the histogram of the true distribution, *h*(*p*), rounded so as to be supported on the set *X* of discrete probability values, is a point in the *plausible* region of the linear program in UnseenEst2. Intuitively the plausible region captures all the histograms that can plausibly generate the observed SFS. The last component of the proof will argue that any two points in the plausible region must be close in generalized relative earthmover distance. This completes the proof because the solution returned by the linear program in UnseenEst2 is in the plausible region and hence must be close in relative earthmover distance to the rounded true histogram, which is close to the true histogram.

The proof of Theorem 2.1 follows the steps of the proof of Theorem 2 in [7]. We have to replace calculations involving Poisson distributions with Binomials in the appropriate places. We also have to rescale all the earthmoving costs by ∑ *p_s_*. We provide explicit analysis where our proof differs from that of [7]; otherwise we refer to the appropriate part of [7] when the calculations are identical.

#### 6.1 Faithful samples

##### Definition 6.1.

*A sample of k alleles with fingerprint* 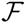*, drawn from a set P* = {*p_s_*} *of probabilities with histogram h and sum t* = ∑*_s_ p_s_, is said to be* faithful *if the following conditions hold:*

- |*m* − *kt*| ≤ (*kt*)^0.6^.
- *For all i,*

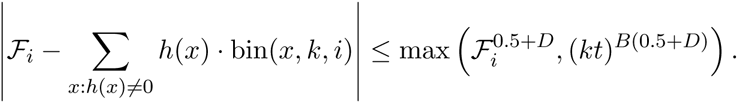
- *For all possible variants* 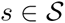, *letting p_s_ denote the true probability of *s*, the number of times *s* occurs in the sample from P differs from its expectation* 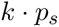 *by at most*

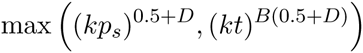

##### Lemma 6.2

(Analogous to Lemma 11 in [7]). *There is a constant γ* > 0 *such that for sufficiently large number of individuals, k, the empirical distribution is* faithful *with probability at least* 1 − *e^−^*^(^*^kt^*^)^*^γ^, where t* = ∑*_s_ P_s_.*

###### Proof.

The first condition follows from Hoeffding bound with high probability.

In our model, 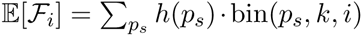. Each fingerprint 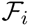 is the sum of independent binary variables, representing whether each mutation occurred exactly i times in the population. Hence Chernoff bounds apply. The analysis showing that the second condition is satisfied is the same as in the proof of Lemma 11 in [7]. We include it here for completeness.

The analysis of the second condition is split into two cases, according to whether 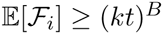. If 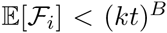, we have that 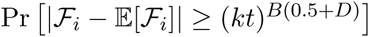 is upper bounded by the case where 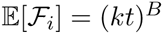. By Chernoff bound,

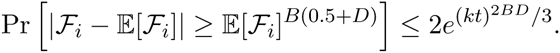

In the case that 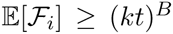, we have that 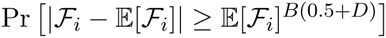 is monotonically decreasing in 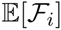 and hence this quantity is bounded by setting 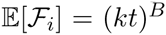. A union bound over the first 2*kt* fingerprints shows that the probability that a sample of *k* alleles violate the first condition is at most 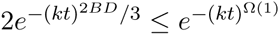. Note that the probability that there are more than 2*kt* nonzero fingerprints is similarly bounded, as the probability that a variant is observed more than 2*kt* times is inverse exponential in *kt*.

For the third condition, we want to show that for all variants *s*, the number of times that *s* is observed in *k* alleles differs from its expectation *p_s_k* by at most max((*kp_s_*)^0.5+D^, (*kt*)^B(0.5+^*^D^*^)^). The analysis also splits into two cases depending on whether *p_s_k* ≥ (*kt*)*^B^* and follows from the same Chernoff bound as before, replacing 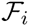 by the number of times *s* occurs in the sample and replacing 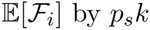 by *p_s_k*. □

##### Definition 6.3.

*Given a fingerprint* 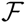*, an upper bound on the support size n,* 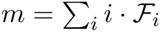*, and a finite set of probability values X, the plausible region is the set of histograms h supported on X satisfying the conditions*

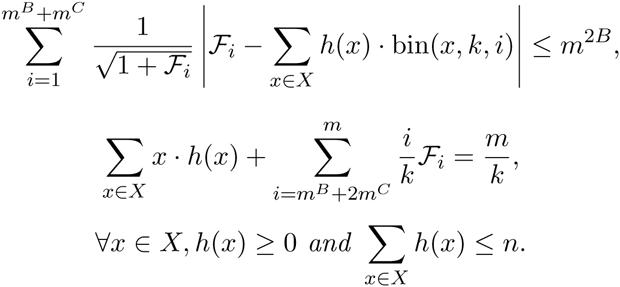

As the name suggests, the plausible region is the set of histograms that can plausibly generate the observed fingerprint 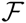. The last three requirements of plausibility are the same as the LP constraints in UnseenEst2.

The following lemma shows that, given a faithful sample of *k* alleles, the corresponding plausible region has a point that is extremely close to the histogram of the true distribution.

##### Lemma 6.4.

*(Analogous to Lemma 12 of the [7].) For sufficiently large k, and n < m*^2+B/2^/*k: given a distribution of support size at most n and a* faithful *sample of k alleles with fingerprint* 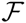*, the plausible region has a point v*′ *such that v′ is close to the true histogram h*

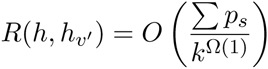

*where h_v_*_′_ *is obtained from* v*′ by appending the empirical fingerprint entries* 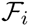 *for i ≥ m^B^* + 2*m^C^.*

###### Proof.

The idea of the proof is to show that, provided the sample is faithful, the true histogram *h* can be minimally modified into a plausible point *v′*. We construct *v′* by taking the portion of *h* with probabilities at most 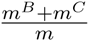 and rounding the support of *h* to the closest multiple of 1/*m*^2^, so as to be supported at points in the set *X* = {1/*m*^2^, 2/*m*^2^,…}.

We construct *h*′ and *v′* as in [7]. The first two steps of the construction are the same. In the third step, we want to normalize the total probability mass 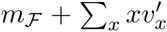 to be *m/k* instead of to 1. This involves rescaling *v′_x_* by a factor of 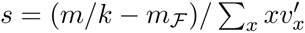.

Next we show that; the discretization does not violate the requirements of plausibility. We note that 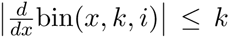. Since we discretize to multiples of 1/*m*^2^, the discretization alters the contribution of each site to each expected fingerprint by at most *k/m*^2^. The support size is bounded by *n*, the discretization alters each expected fingerprint by at most *n* · *k/m*^2^. The rescaling step also does not violate the plausibility conditions. Finally the last part of the proof bounds the per unit earth-moving cost, which does not use any properties of the Poisson distribution. We can apply the same earth-moving scheme and analysis of the per unit cost. The final cost *R*(*h*, *h_v′_*) needs to be scaled by *m/k* since that’s the total amount of probability mass. □

#### 6.2 Chebyshev construction

The previous section established that, given a faithful sample (which we are likely to obtain with high probability), there exists a plausible point which is very close to the true histogram. In this section, we will show that any two plausible points are close in generalized relative earthmover distance. By the triangle inequality, this guarantees that the solution returned by UnseenEst2 will be close to the true histogram. To establish the closeness of the histograms, we will explicitly construct a earthmoving scheme using Chebyshev polynomials. This is analogous to the earthmoving scheme in [7], replacing all instances of poi(*kx*, *i*) by bin(*x, k, i*).

##### Definition 6.5.

*For a given k, a β-bump earthmoving scheme is defined by a sequence of positive real numbers* {*c_i_*}, *the bump centers, and a sequence of functions* 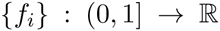 *such that* ∑*_i_ f_i_*(*x*) = 1 *for each x and each function f_i_ may be expressed as a linear combination of Binomials, f_i_*(*x*) = ∑*_j_ a_ij_*bin(*x*, *k*, *j*) *such that* ∑*|a_ij_| ≤ β. Given a generalized histogram h, the scheme works as follows: for each x such that h*(*x*) ≠ 0, *and each integer i* ≥ 0, *move xh*(*x*) · *f_i_*(*x*) *units of probability mass from x to c_i_. We denote the resulting histogram by* (*c, f*)(*h*).

We define binomial Chebyshev bumps, following [7].

##### Definition 6.6.

*Let s* = ⌊0.2 log *kt*⌋, *where t* = ∑*_s_p_s_. Define* 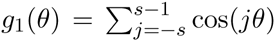 *to be an approximation of the delta function, truncated at Fourier degree s. Define a slightly fatter version*

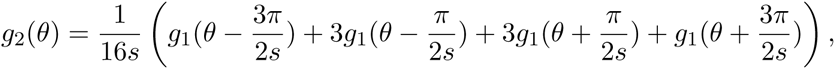

*and, for i* ∈ {1,…,*s* − 1}, *define its shifted versions* 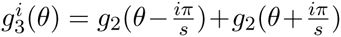*, and* 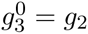*, and* 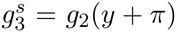*. Let t_i_*(*x*) *be the linear combination of Cheybyshev polynomials so that* 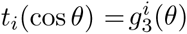. *We define s* + 1 *functions, the “skinny bumps”, to be* 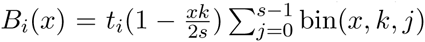*, for i* ∈ {0, …, *s*}.

##### Definition 6.7.

*The Chebyshev earthmoving scheme is defined in terms of k as follows: let s* = 0.2 log *kt. For i ≥ s* + 1, *define the ith bump function f_i_*(*x*) = bin(*x*, *k*, *i*) *and associated bump center* 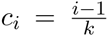*. For i* ∈ {0, …,*s*} *let f_i_*(*x*) = *B_i_*(*x*) *and define their associated bump centers* 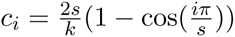*, and let c*_0_ = *c*_1_.

We now prove a number of nice properties about the Chebyshev earthmoving scheme.

##### Lemma 6.8

(Lemma 18 in [7]). *For any θ,*

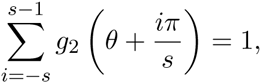

*and for any x,*

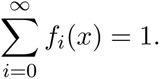

###### Proof.

Same as in Lemma 18 of [7]. Nothing special about Poisson density was used in that proof. □

##### Lemma 6.9

(Analogous to Lemma 19 in [7]). *Each B_i_*(*x*) *may be expressed* as 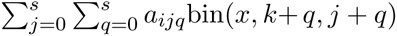 *for a_ijq_ satisfying*

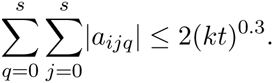

###### Proof.

We decompose 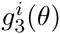 into a linear combination of cos(*ℓθ*), for *ℓ* ∈ {0,…, *s*}. Since cos(−*ℓθ*) = cos(*ℓθ*), *g*_1_(*θ*) consits of one copy of cos(*sθ*), two copies of cos(*ℓθ*) for each *ℓ* strictly between 0 and *s*, and one copy of cos(0*θ*). *g*_2_(*θ*) consists of (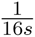 times) 8 shifted copies of *g*_1_(*θ*)’s. The shifts changes the phases of the Fourier coefficients but not their magnitude. Sine components may have been introduced in the shifts, but since 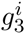 is an even function, the sine components cancel out. Since each *g*_3_ contains at most two shifted *g*_2_’s, each 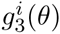 is a linear combination 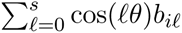 with the Fourier coefficients bounded by 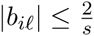.

Since *t_i_* was defined so that 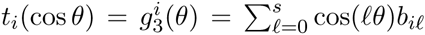, by the definition of Chebyshev polynomials we have 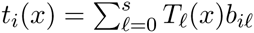. Thus the bumps are expressed as

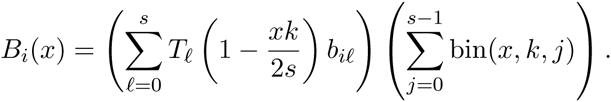

We further express each Chebyshev polynomial via its coefficients as 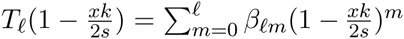. We then expand each term via binomial expansion as 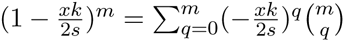 to yield

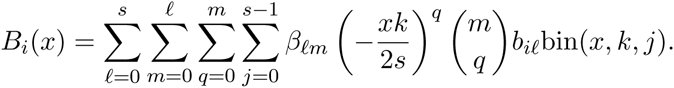

In general we can re-express

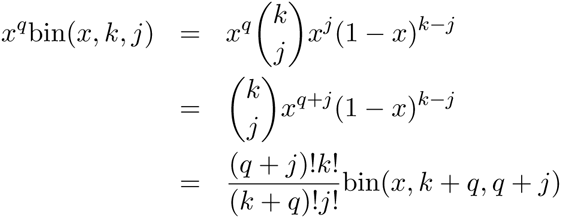

Following the same calculations as in the Unseen, we have

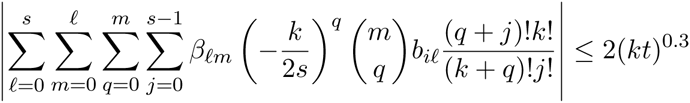

##### Lemma 6.10

(Lemma 20 in [7]). 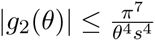 *for θ* ∈ [−*π*, *π*] \ (−3*π*/*s*, 3*π*/*s*), *and* |*g*_2_(*θ*)| ≤ 1/2 *everywhere.*

###### Proof.

Same proof as in Lemma 20. This lemma doesn’t involve Poisson density at all. □

##### Lemma 6.11

(Analogous to Lemma 21 in [7]). *The Chebyshev earthmoving scheme is* 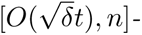 *good*, *where* 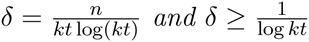 *and* 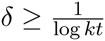.

###### Proof.

The analysis has two parts. For the first part, we consider the cost of bumps *f_i_* for *i* ≥ *s* +1, where recall that *s* = 0.2 log *kt*. This is the cost of moving bin(*x, k, i*) mass from *x* to 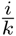. The unit cost of moving mass from *x* to 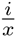 is 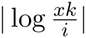, which is upper bounded by 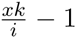 when *i* < *xk* and 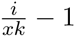 otherwise. We split the calculation into two parts. First, for *i* > ⌈*xk*⌉,

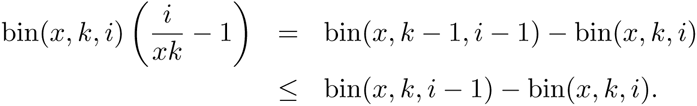

When summed over *i* ≥ max{*s*, ⌈*xk*⌉}, this telescopes to an expression bounded by

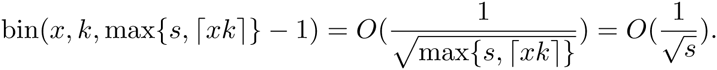

For *i* ≤ ⌈*xk*⌉ − 1, since *i* ≥ *s*, we have 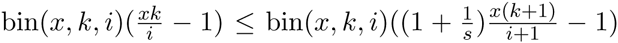. The 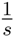 term sums to at most 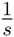. Note that 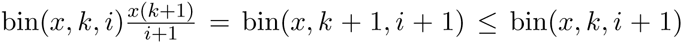, where the last inequality is because *i* < ⌈*xk*⌉ − 1. Therefore the rest of the sum telescopes to 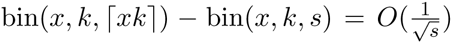. Thus in total, *f_i_* for *i* ≥ *s* + 1 contributes 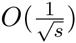 to the relative earthmover cost, per unit of weight moved.

Next we analyze the skinny bumps *f_i_*(*x*) for *i* ≤ *s*. The simple case is when *xk* ≥ 4*s*. Recall the definition 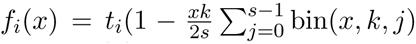. Since *xk* > *x*, we bound 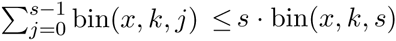. Each *f_i_*(*x*) is exponentially small in both *x* and *s*, the thus the total earthmoving scheme, per unit of mass above 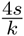 is exponentially small.

The remaining case is *xk* ≤ 4*s* and *i* ≤ *s*. The trigonometric calculations here does not use any properties of Poisson distributions and carry over without change to our Binomial case. The per unit earthmoving cost in this regime is 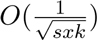. For a distribution with histogram *h*, the cost of moving earth on this region, for bumps *f_i_* where *i* ≤ *s* is thus

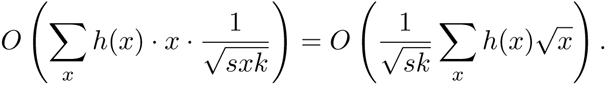

Since ∑*_x_ x* · *h*(*x*) = *m/k* and ∑*_x_ h*(*x*) ≤ *n*, by the Cauchy-Schwarz inequality,

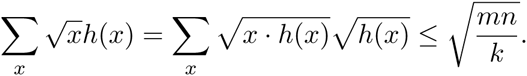

The total earthmoving cost in this regime is 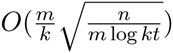 and hence we need *n* = *δm* log *kt* to ensure that the total cost here is 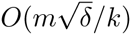.

Finally we put all the pieces together. The total probability mass that need to be moved is *O*(*m/k*). The regimes of *i* ≥ *s* + 1 and *i* ≤ *s*, *xk* ≥ 4*s* both require 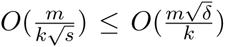 earthmoving cost, since *s* = 0.2 log *kt* and 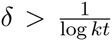 by assumption. The last regime of *i* ≤ *s*, *xk* ≤ 4*s* also incurs 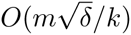 cost and hence the overall earthmoving cost is 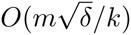. □

###### Proof of Theorem 2.1.

To wrap up the proof of the theorem, let *g* be the generalized histogram returned by the linear program and let *h* be the plausible point constructed to be close to the true histogram *p*, 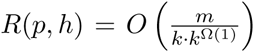. Let *h*′ and *g*′ be the generalized histograms that result from applying the Chebyshev earthmoving scheme to *h* and *g*, respectively. We have 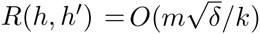 and 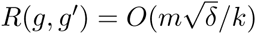.

What is left if to bound *R*(*g*′, *h*′) by 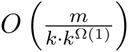. For the bump centers *i* ≥ *s* + 1, the same analysis as in [7] shows that relative earth mover cost is 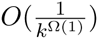. We consider teh first *s* + 1 = *O*(*logkt*) bump centers corresponding to the skinny Chebyshev bumps. Recall that for these centers, *c_i_*, the bump functions *B_i_*(*x*) may be expressed as 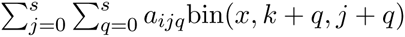 for *a_ijq_* satisfying

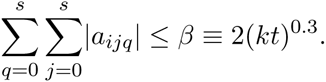

Using the shorthand ∑*_x_* for ∑*_x:h_*_(_*_x_*_)+_*_g_*_(_*_x_*_)≠0_, we have

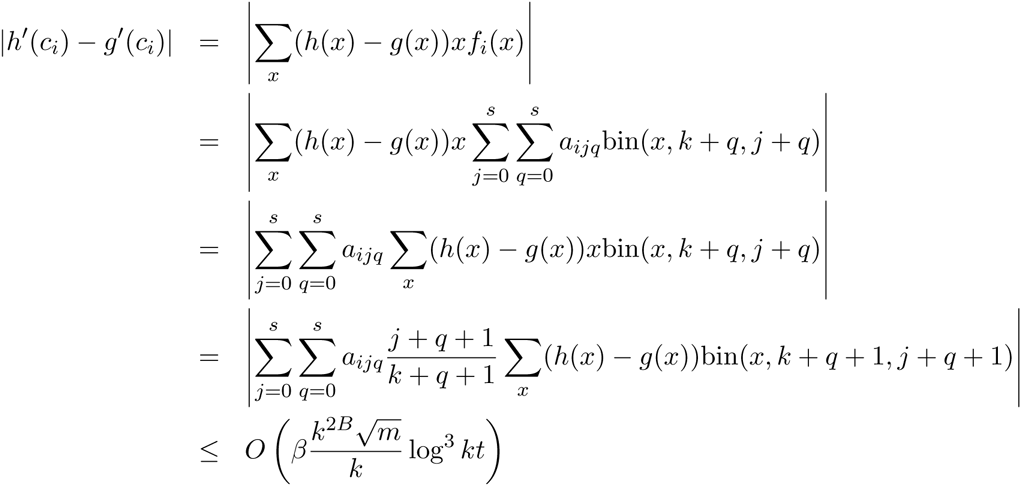

where we have used triangle inequality and the first condition of plausibility in the last inequality. Since *B* < 0.1, we have that this discrepancy is 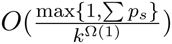 for each center *c_i_*, and since there are log *kt* centers, the total discrepancy is also 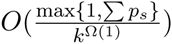. Putting all the pieces together, by the triangle inequality, we have

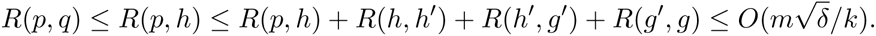

Moreover, 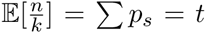 and since alleles are independent, Chernoff bounds applies and with probability at least 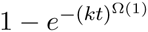, 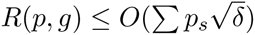. □

#### 6.3 Proof of Proposition 1.4

For convenience, we restate the proposition in a slightly more general form:

##### Proposition 1.4

*Given two lists of probabilities* 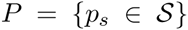 *and* 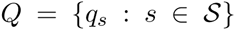 *with ∑_s_ P_s_* > ∑*_s_ q_s_, let* 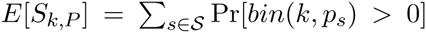 *denote the expected number of variants observed in a sample of k alleles with the distribution of frequencies given by P, and let E*[*S_k_,_Q_*] *denote the analogous quantity corresponding to frequencies Q. Let* 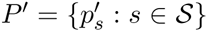 *be any list of probabilities satisfying:*

1. *Either for all* 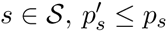, *p*′*_s_* ≤ *p_s_*, *or for all* 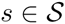, *p*′*_s_* ≥ *p_s_*,
2. ∑*_i_ p*′*_s_* = ∑*_i_ q*_s_,

*then, for any k,*

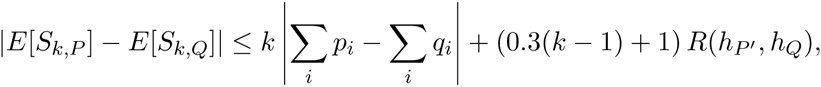

*where R*(*h_P_*_′_, *h_Q_*) *is the relative earthmover distance between the histograms corresponding to P′ and Q. Hence for k* > 3,

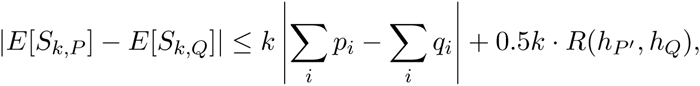

###### Proof.

By the triangle inequality, |*E*[*S_k,P_*] − *E*[*S_k,Q_*]| ≤ |*E*[*S_k,P_*] − *E*[*S_k,P_*_′_]| + |*E*[*S_k_*,*_P_*_′_] − *E*[*S_k,Q_*]|. The first term is trivially bounded by *k* ∑*_i_* |*p_i_* —*p′_i_* | = *k* |∑*_i_ p_i_* − ∑*_i_ q_i_*|, since each unit of probability mass can, in expectation, account for at most *k* distinct observations. To bound the second term, first note that both the relative earthmover cost, and expected number of distinct elements observed are linear functions of the number of elements of *P* and *Q* with each different probability value, it suffices to analyze the costs of the earthmoving distance and the change in the expected number of distinct elements for a single earthmoving operation: consider moving *c* units of mass from probability value *x* to *y*. The change to the expected number of distinct elements observed is exactly

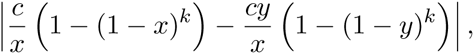

and the relative earthmover cost of this is 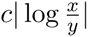. We now show that the ratio of these quantities is always at most 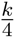.

We seek to bound the maximum change in 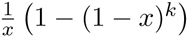 relative to the change in log *x* as *x* changes, namely the maximum ratio of their derivatives, where we add a negative sign since 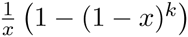 is a decreasing function. Since 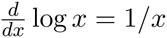, the ratio of derivatives is

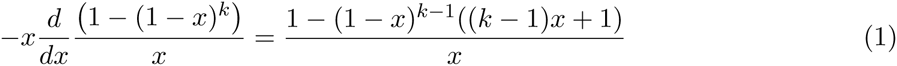

Consider the approximation (1 − *x*)*^k^*^−1^ ≈ *e* ^−^*^x^*^(^*^k^*^−1)^. Taking logarithms of both sides, and using the fact that 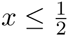 we have log 1 − *x* ≥ − *x* − *x*^2^, we have that for 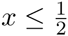 the inequality (*k* − 1) log(1 − *x*) ≥ − (*k* − 1)(*x* + *x*^2^); exponentiating yields 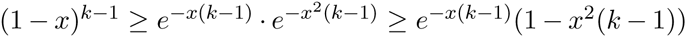.

Thus for 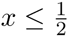 the ratio of derivatives is bounded as

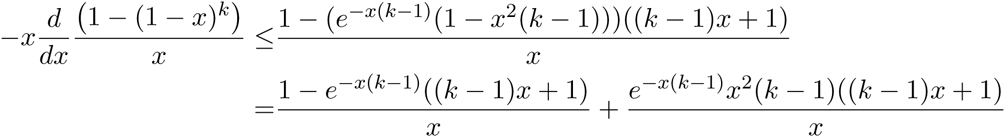

The first term of the right hand side, after dividing by *k* − 1, can be reexpressed in terms of *y* = *x*(*k* − 1) as 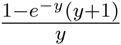, which has a global maximum less then 0.3; the second term in the right hand side, after the same variable substitution, equals *e*^−^*^y^y*(*y* + 1), which has a global maximum less than 1. Thus, for 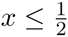, the absolute value of the ratio of derivatives is bounded as 0.3(*k* − 1) + 1. For 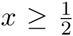, the right hand side of Equation 1 is 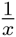 minus some positive quantity, and is hence at most 2. Since 0.3(*k* − 1) + 1 ≥ 2 for any *k* ≥ 5, all that remains is to checking the *k* = 2,3,4 cases where 0.3(*k* − 1) + 1 < 2 by hand to confirms that 0.3(*k* − 1) + 1 is in fact a global bound. □

